# Ethanol down-regulates gastrula gene expression and cell movement, causing symptoms of foetal alcohol spectrum disorders

**DOI:** 10.1101/2024.11.30.626151

**Authors:** Amena Ali Alsakran, Hoi Ying Wong, Caitlin Heaton, Rebekah Boreham, Jonathan Ball, Tetsuhiro Kudoh

**Affiliations:** Biosciences, University of Exeter, Exeter, EX4 4QD, United Kingdom

## Abstract

Foetal alcohol spectrum disorders (FASDs) occur in embryos when they are exposed to maternally supplied alcohol. To study the mechanisms of FASDs, the zebrafish embryo can serve as an excellent model as ethanol exposed zebrafish embryos exhibit common symptoms of human FASDs including microcephaly, incomplete neural plate closure, eye defects, craniofacial disorders and many other defects. Here we investigated the embryo development at gastrula stage when three germ layers develop with specific gene expressions and undergo dynamic cell movement including extension, convergence and epiboly, establishing the platform to form the head and body axis in the later development. Gastrula cell movement analyses using fluorescent transgenic zebrafish embryos revealed that ethanol induced dose dependent delay of extension, convergence and epiboly cell movement and associated gene expressions in all three germ layers. Our results suggest multiple targets of ethanol including gene expression and cell movement, consequently delay the key gene expression and cell localisation, causing irreversible developmental defects in the head and body axis formation.

## Introduction

Foetal alcohol spectrum disorders (FASD), is a term used to represent a spectrum of birth issues due to mothers’ alcohol consumption. Approximately 9.8% of pregnant women worldwide consume ethanol, with a reported prevalence of FASD of 4.4 per 1000 among children born in the United States (Popova *et al*., 2019). Multiple FASD-related abnormalities can occur in the fetus’s central nervous system (Wilhelm 2016), such as microcephaly, incomplete neural tube closure, mental retardation and blindness with suppressed retinal development. (O’Neil, 2014).

FASD research largely relies on translational animal models, such as rodents and zebrafish, to discover the mechanisms of toxicity of ethanol (EtOH) in embryonic and foetal development (Pinheiro-da-Silva & Luchiari, 2021). Zebrafish embryo has been an ideal model for investigating various angles of ethanol toxicity, including cellular and genetic toxicity and toxic responses at the organism level (Alsakran and Kudoh, 2021). Features such as embryo transparency, time-efficient analyses, high fecundity, short life span, and toxicity specific biosensor transgenic technology, availability of genetic data favour their application in toxicity assessment studies (Modarresi et al., 2020, Lee et al. 2012, Mourabit et al. 2019). Ethanol-exposed zebrafish embryos exhibit pre-hatching and post-hatching growth deficiencies, which are similar to FASDs in children (Arenzana *et al*., 2006). Ethanol negatively affects various aspects of the zebrafish life cycle in a dose-dependent manner, which include disturbed gastrulation cell movement, embryonic development, cell death, alteration in stem cell gene expression, microcephaly, CNS morphogenesis, neuronal development and skeletal dysmorphogenesis (Alsakran and Kudoh, 2021; Carvan *et al*., 2004). Retinal cell differentiation and proliferation were also affected in response to ethanol exposure (Fu et al., 2021). Multiple studies have reported ethanol toxicity-based defects in the early stages, which include delayed epiboly resulting from abnormal cell adhesion and movement (Sarmah 2013, Alsakran and Kudoh, 2021). Marrs *et al*., (2010) reported a reduction in embryo body length in addition to epiboly delay, while Sarmah *et al*., (2020) revealed the disruption of the microtubule cytoskeleton that restricts the formation of microtubule filament, which is crucial for epiboly movement during gastrulation. Although cell movement and gene expression are both affected by ethanol at the gastrula stage, it remains unknown if these effects are mutually linked with an epistatic relationship. It is also unclear how these early defects at gastrula specifically cause later morphological abnormalities such as a small brain, small eyes, open brain and other morphological abnormalities. In this project, we analysed gastrula cell movement in detail using a large number of samples the ACQUIFER multiwell time-lapse imaging system using fluorescent transgenic zebrafish lines Tg(h2a:gfp) and Tg(gsc.gfp) which are suitable for visualising the morphology of blastoderm and axial mesoderm respectively. We have also analysed key gene expression at the gastrula stage in all three germ layers, endoderm, mesoderm and ectoderm that have a crucial role in the following embryonic morphogenesis. By combining these results, we will discuss the toxicity mechanisms of EtOH with linking the gastrula stage cell movement, gene expression and following development and body patterning.

## Materials and Methods

### Zebrafish (*Danio rerio*)

Zebrafish strains, Wild Indian Karyotype (WIK), Tg(h2a:GFP) and Tg(gsc:GFP) lines were maintained at the University of Exeter’s Aquatic Resource Centre (ARC) supplied with aerated and circulated freshwater dissolved with artificial sea salt (0.2ppt), 28°C ± 1°C. Zebrafish eggs were collected by natural spawning in the morning within 30min after fertilisation. The embryos in petridishes were incubated at 28 °C.

### Ethanol treatment

Zebrafish embryos were treated with different concentrations of ethanol (0.5%, 1%, 2% and 3%) starting from the 2hpf either in 10cm petridish for live imaging or sampling for staining or in 96-well plate for ACQUIFER automated time-lapse imaging. Live images of embryos were captured using Nikon SMZ1500 stereomicroscope. The image was analysed by measuring epiboly and extension of the axial mesoderm. Embryos were randomly oriented in 96 well plates, therefore the progress of epiboly was calculated by estimating the distance between the animal pole and the plane of the blastoderm margin. The statistical difference among treatments was established by subjecting the data to one-way ANOVA, and means were compared using Tukey’s multiple comparison test at p<0.0001.

### Automated time lapse imaging and analyses of gastrula cell movement

Control and EtOH treated zebrafish embryos with transgenic backgrounds, Tg(h2a:gfp) or Tg(gsc:gfp) were incubated in a 96 well plate at 28°C from 4hpf to the end of gastrulation. During the period, both transmission light and fluorescent images with GFP filter (488nm) were captured every hour. Using FIJI ImageJ, progress of elongation of blastoderm margin and extension of the axial mesoderm were measured. From the image of Tg(h2a:gfp) embryos, oval of the blastoderm margin was identified, and the distance between the centre of the oval and the animal pole was measured to calculate the length of the blastoderm. As for the extension of the axial mesoderm, length of the GFP-positive axial mesoderm structure in the Tg(gsc:gfp) embryos were measured.

### *In situ* hybridisation

Zebrafish embryos were fixed with 4% PFA in PBS with chorion and manually dechorionated in PBS with tweezers. Subsequently embryos were stored in methanol at -20’C. Preparation of probes and in situ staining methods in zebrafish are previously reported (Kudoh et al., 2004).

## Ethical declaration

All experimental methods were approved by the University of Exeter ethical committee, UK Home Office animal project license and carried out in accordance with these guidelines.

## Results

### Live imaging of zebrafish WT embryos treated with EtOH reveals dose dependent morphological abnormalities and lethality

To examine the effect of alcohol at early embryonic development around the blastula to gastrula stage, zebrafish embryos were treated with ethanol from 2hpf (early blastula) with a series of concentrations from 0.5% to 3%. Higher mortality rates were observed with increasing concentrations of EtOH during blastula, gastrula and following segmentation stages (Figure 1). The highest mortality rates were recorded with 3% ethanol showing a 40% mortality rate for blastula to gastrula and a 66.7% rate for the somitogenesis stages. A 100% survival rate was noted with 0% and 0.5% concentrations for all embryonic stages. Even at the lowest dose of 0.5% EtOH, a degree of deformity was observed at the segmentation stages in the form of oedema in the heart cavity (23.3%), which was also unequivocally noted at 1% and 2% EtOH.

**Figure 1.**
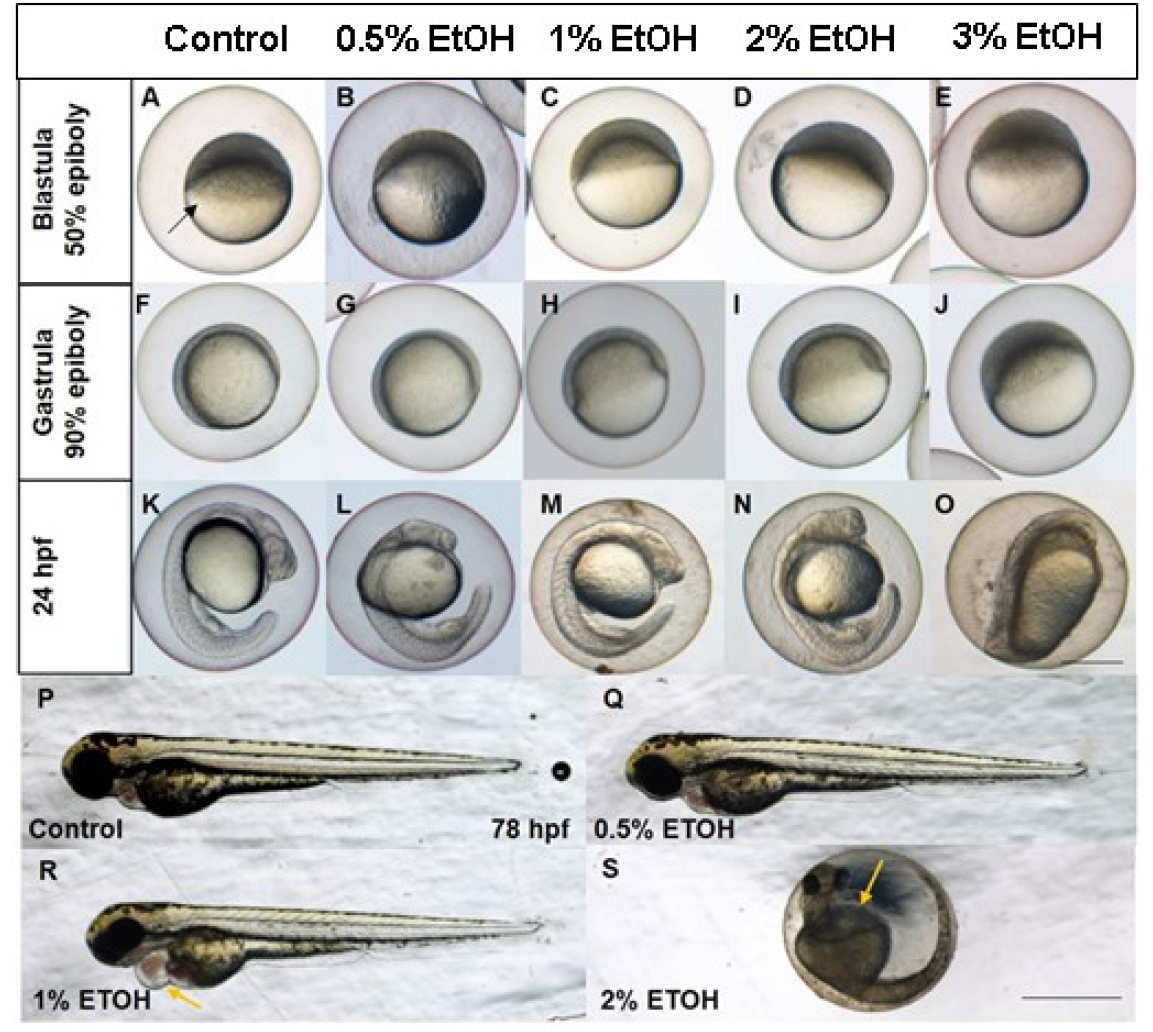
Morphological defects observed in the ethanol treated embryos at late blastula (A-E), late gastrula (F-J), 24hpf (K-O) and 78hpf (P-S). Scale bar = 500um (A-O) or 2mm (R-S).

### Fluorescent transgenic zebrafish embryos treated with EtOH show dose dependent delay of epiboly at gastrula stage

The transgenic line, Tg(h2a:gfp) show broadly expressed green fluorescence in the blastoderm and is suitable for visualising and measuring the progress of epiboly cell movement at gastrula stage. To examine the level of epiboly delay caused by ethanol, we used this line to observe the (Figure 2). There was a noticeable delay in the epiboly movement with EtOH groups in a dose dependent manner. The mean epiboly (%) values at 5hpf remained significantly different from each other (37%, 33% & 34% at 1%, 2% and 3% EtOH concentrations) and from the control treatments (47% at 0% EtOH). There was a gradual delay of epiboly from 5 hpf until 9 hpf. The mean epiboly value of the control (0% EtOH) at 9hpf was (92%) compared with EtOH treated embryos (73%, 56% & 42% at 1%, 2%, and 3% ETOH concentrations, respectively) (Figure 2).

**Figure 2.**
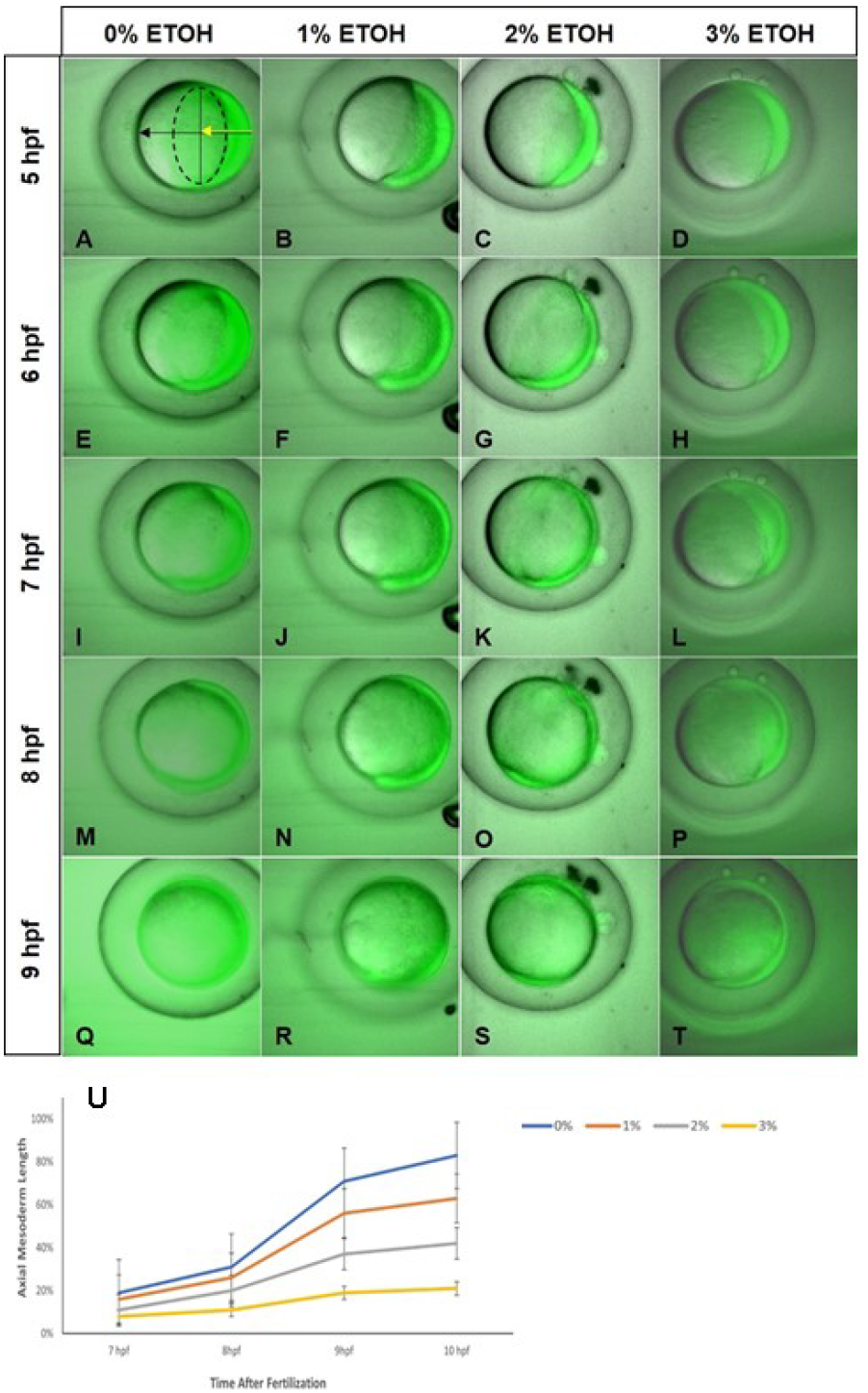
Tg(h2a:gfp) zebrafish embryos reveal dose dependent delay of epiboly. The Tg(h2a:gfp) line shows expansion of the blastoderm during gastrulation. A-D 6hpf, E-H 7hpf, I-L 8hpf, M-P 8hpf and Q-T 9hpf. U. shows measurement of progress of epiboly in the control and ethanol treated embryos revealing dose dependent delay of epiboly by ethanol from 0%, 1%, 2% to 3%.

### TG (gsc:GFP) zebrafish embryos treated with EtOH reveal delay of extension cell movement at gastrula stage

The Tg(gsc:gfp) line markes the axial mesoderm (prechordal plate & notochord) with GFP and is suitable for visualising the extension movement of the axial mesoderm (Boutillon et al., 2022) (Figure 3). The data shows that the axial mesoderm extension is delayed by EtOH exposure. At the end of our measurement, the mean growth values at 10hpf were also found to be significantly different from each other (83%, 63%, 42%, 21% at 0%, 1%, 2%, and 3% ETOH concentrations) (Figure 3). These data suggest that EtOH also delay the extension cell movement during the gastrula stage.

**Figure 3.**
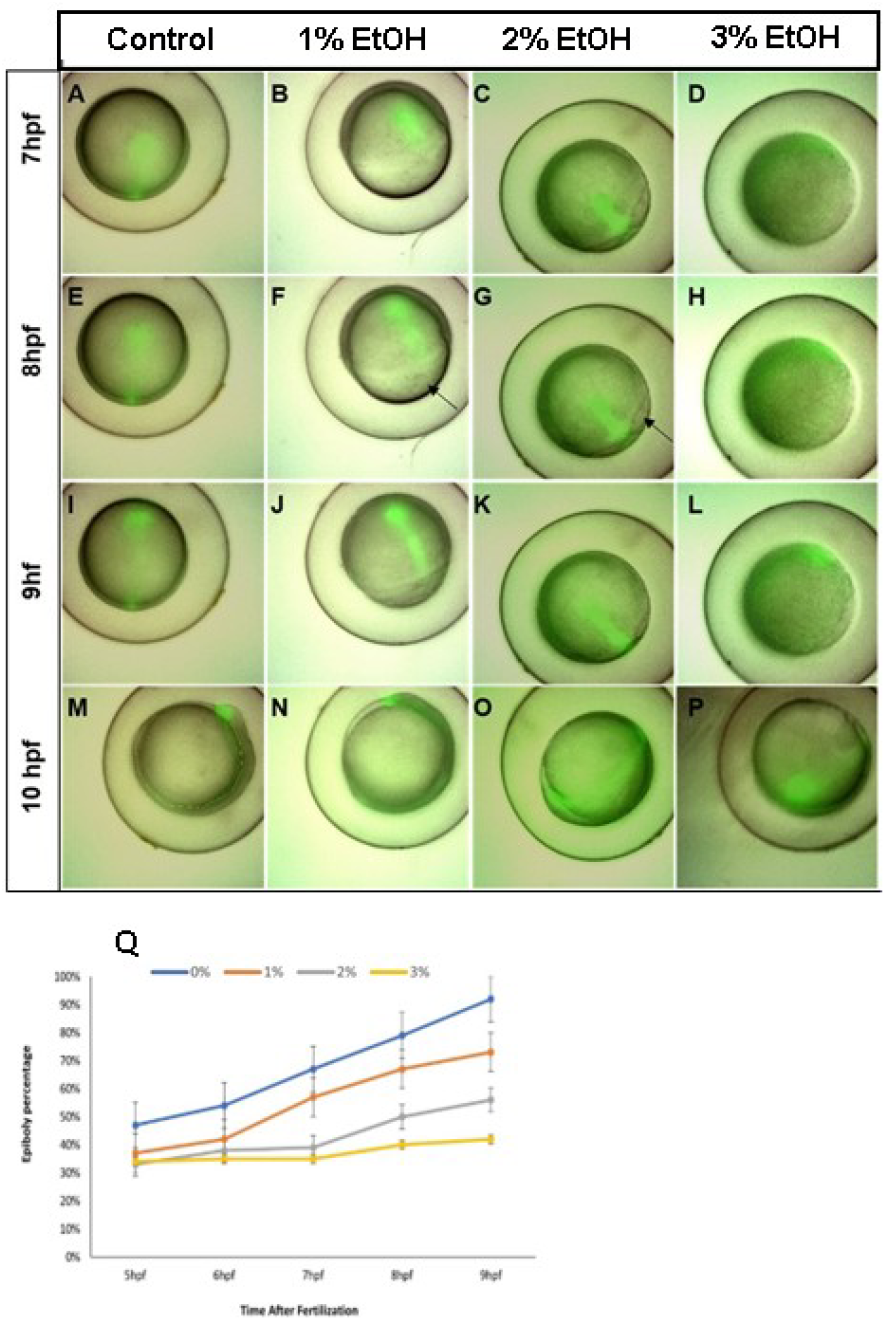
Tg(*gsc*:gfp) zebrafish embryos reveal dose dependent delay of epiboly. The Tg(*gsc*:gfp) line shows expansion of the blastoderm during gastrulation. A-D 7hpf, E-H 8hpf, I-L 9hpf, M-P 10hpf. Q. shows measurement of progress of epiboly in the control and ethanol treated embryos revealing dose dependent delay of epiboly by ethanol from 0%, 1%, 2% to 3%.

### Gene expression analyses using in situ hybridisation reveal reduction of gene expression in a variety of genes and tissues and delay of the convergence-extension cell movement

As embryo morphology changed by EtOH exposure, expression of marker genes for each embryonic domain and cell lineage at gastrula stage were examined using in situ hybridisation. Figure 4 and 5 show results of markers for endoderm/mesoderm and ectoderm respectively. In the endoderm/mesoderm markers, there are overall trend that gene expression was not clearly suppressed by 1% EtOH but delay of epiboly is visualised by these markers. In addition, convergent-extension movement of endoderm and axial mesoderm cells were reduced (*Sox17*, Fig. 4B and *gsc*, Fig. 4I). In 2% EtOH treated embryos, *sox17, eve1* and *bmp4* were strongly suppressed (Fig. 4C,L,O) but *ntl* and *gsc* were not (Fig. 4I,L).

**Figure 4.**
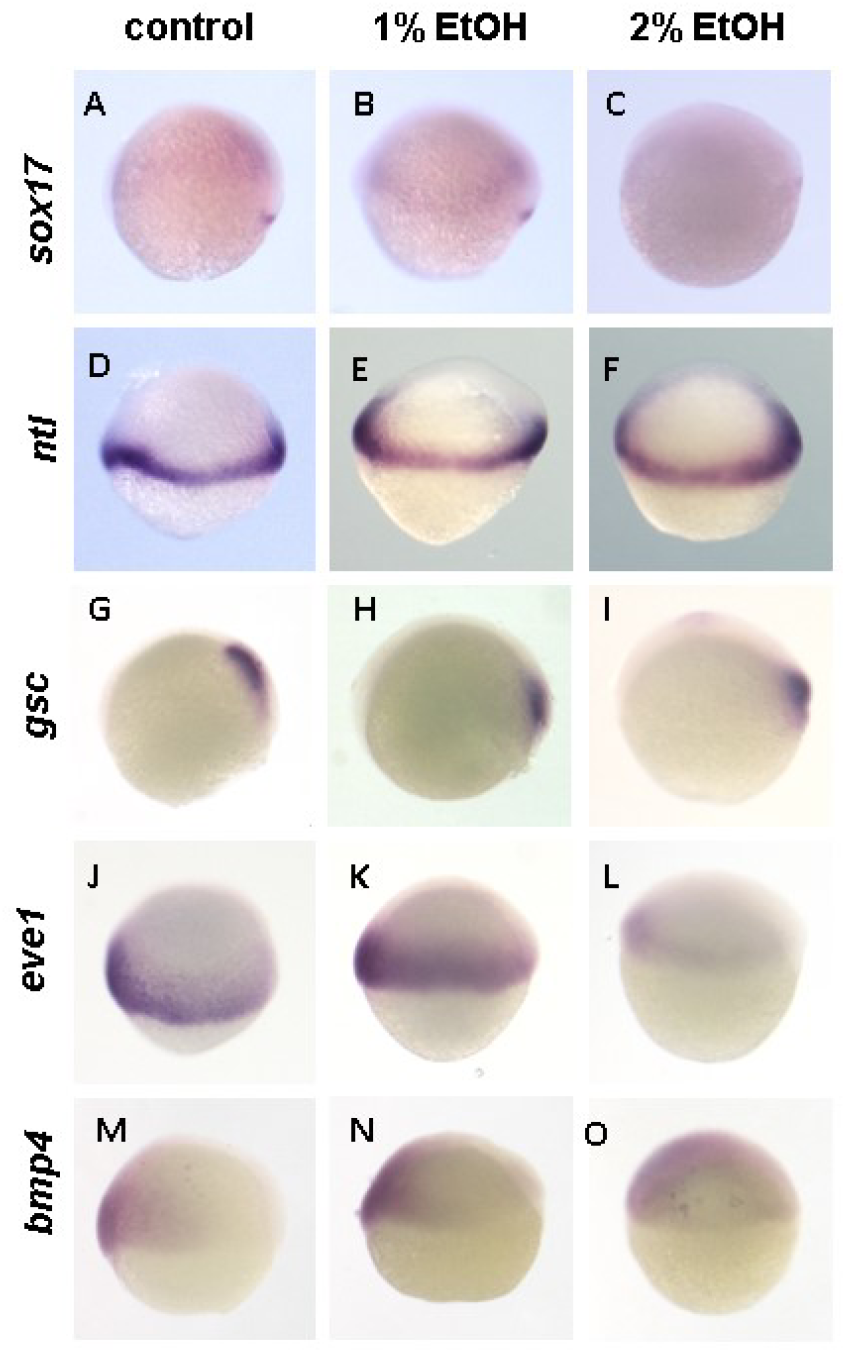
Expression of endoderm and mesoderm genes are suppressed and/or altered by ethanol. In situ hybridisation staining of late gastrula embryos stained with *sox17* (A-C), *ntl* (D-F), *gsc* (G-I), *eve1* (J-L) and *bmp4* (M-O).

**Figure 5.**
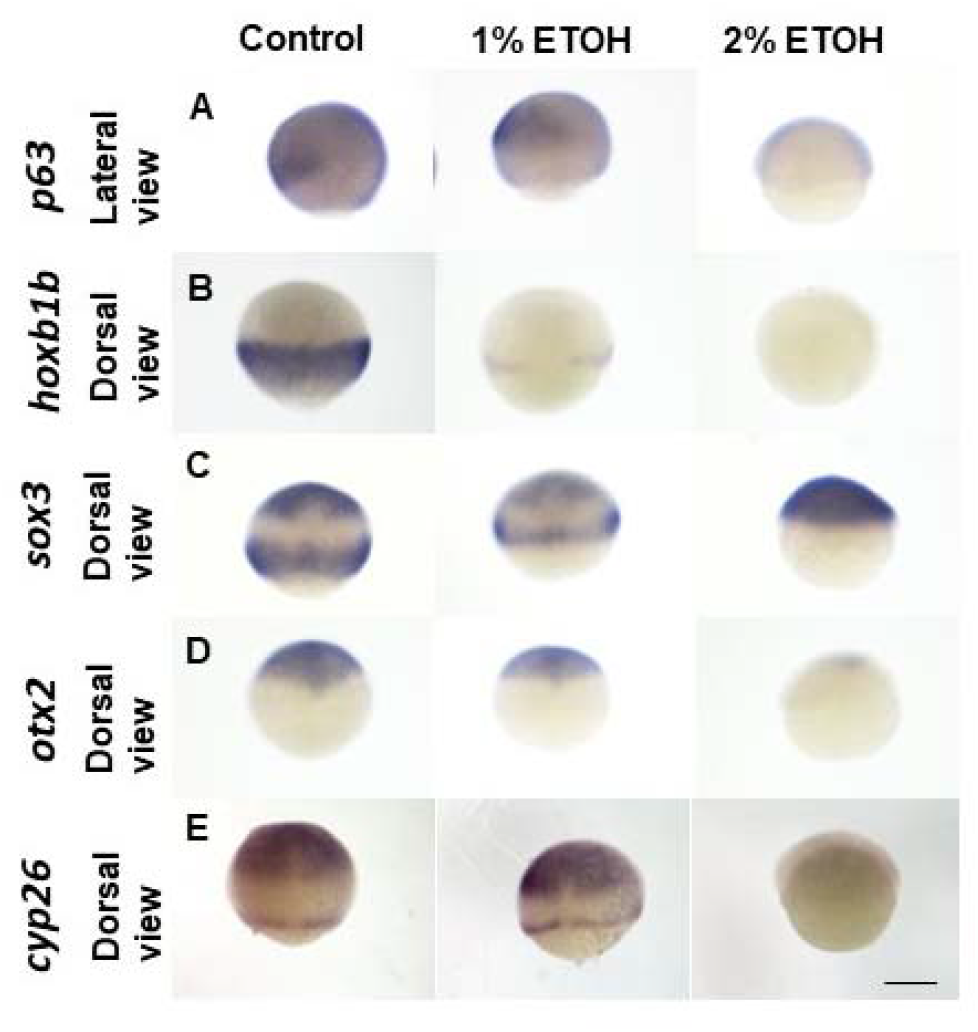
Expression of ectoderm genes are suppressed and/or altered by ethanol. In situ hybridisation staining of late gastrula embryos stained with *p63* (A-C), *hoxb1b* (D-F), *sox3* (G-I), *otx2* (J-L) and *cyp26a1* (M-O).

Consistently to these results, in case of ectoderm markers, most of marker genes were not clearly suppressed with 1% EtOH and only shows narrowing of expression domains due to the epiboly delay (Fig. 5). However, *hoxb1b* showed narrowing of the expression domain with clear decreased of gene expression (Fig. 5E). In 2% EtOH as seen in the endoderm/mesoderm markers, most of ectoderm markers were also strongly suppressed except *sox3* (Fig. 5I). Only the ectoderm marker *sox3* was detected during 2% EtOH exposure of the genes investigated.

Since *sox3* has complicated expression patterns, further analysis of *sox3* expression was conducted with different developmental stages including late blastula, late gastrula and bud stages (Fig. 6). The data revealed that *sox3* has two waves of gene expression, firstly at blastula stage broadly in all ectoderm cells (Fig. 6A) and subsequently in neural ectoderm cells in two domains (anterior and posterior) (Fig. 6D). The stage specific sox3 expression analysis confirmed that 1% EtOH reduce the convergence extension movement of the ectoderm is also suppressed without reducing gene expression of the genes investigated in this study, and with 2% EtOH, gastrula stage specific sox3 expression and gastrula stage specific expression were both slightly reduced.

**Figure 6.**
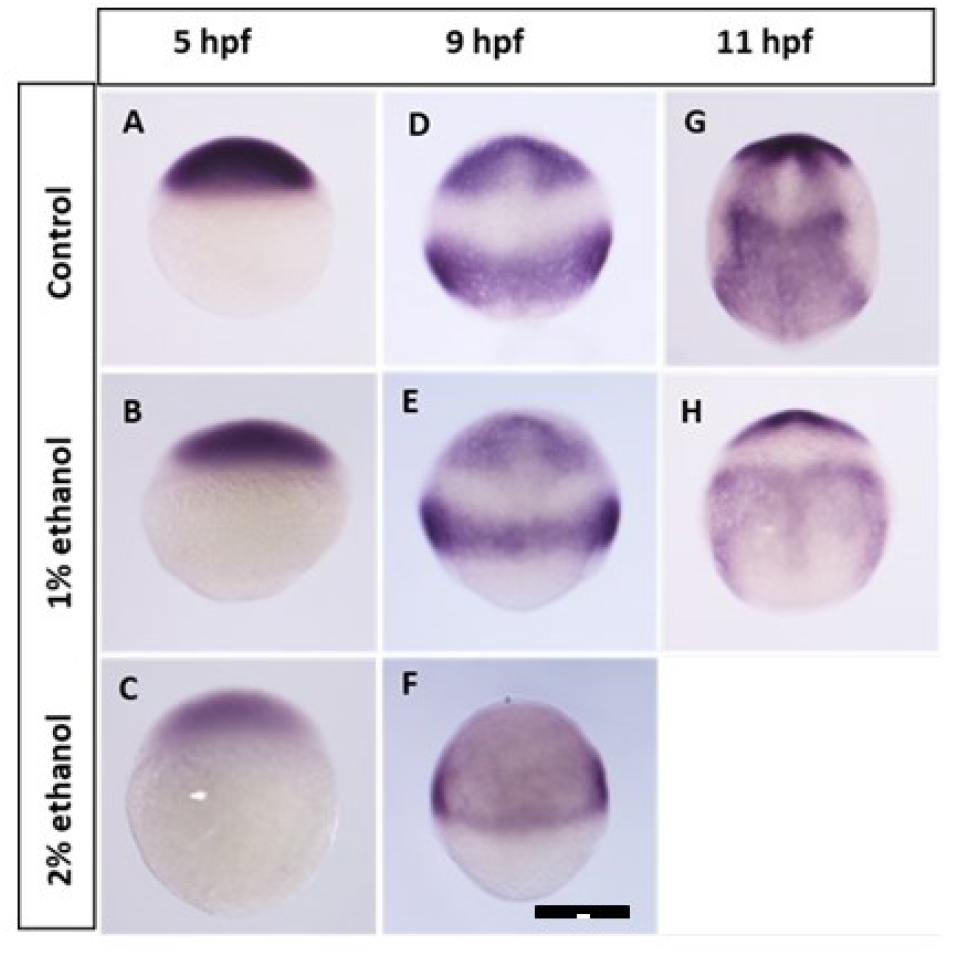
Expression of neural ectoderm gene *sox3* is reduced or delayed by ethanol in a stage specific manner at blastula to gastrula stages. A-C 5hpf, D-F 9hpf, G-I 11hpf. I does not show an image due to high lethality of embryos by this stage.

## Discussion

In this work, we used the zebrafish embryo as a model for studying human fetal alcohol spectrum disorder with focusing on gene expression and cell movement during gastrulation. After ethanol exposure with our condition, microcephaly, reduction of body length and eye size, pericardial edema and enhanced mortality are induced as reported from similar exposure from previous reports (Bilotta et al., 2004; Sylvain et al., 2010, Soares et al., 2012, Joya et al., 2014).

To elucidate the cause of these symptom, we conducted time-lapse live imaging analyses using automated multiwell imaging system for measuring gastrulation cell movement including convergence (cell movement from ventro-lateral toward dorsal), extension (cell movement along animal-vegetal poles) and epiboly (expansion of the blastoderm toward the vegetal pole). To clearly visualise the progress of epiboly, Tg(h2a:gfp), transgenic line which ubiquitously express GFP in the embryonic cells was used (Figure 2). The data showed dose dependent delay or stop of epiboly by EtOH.

The level of delay was partially restored at around bud stage with an intermediate dose of EtOH (e.g. 1%), suggesting that some level of delay can be amended and normal morphological development could still be achieved with a low to medium dose of ethanol at least for the epiboly-mediated developmental processes. However with higher dose of EtOH (2% and 3%) epiboly delay became severer causing incomplete epiboly, failure of the germ ring closure, ceasing the blastoderm to cover the yolk causing embryonic lethality during gastrula stage. Epiboly cell movement is a relatively flexible process in development, and some delay can possibly be restored. For zebrafish, epiboly complete before bud stage, then tail bud is formed and somitogenesis follows. However in medaka, epiboly is complete at the beginning of the somite stage (Iwamatsu 2004). In a more extreme case of the rainbow trout, completion of epiboly delays to the mid-somite stage (Finch and Kudoh 2010). These differences suggest that epiboly delay may not directly affect other essential gross developmental structures such as head development, somitogenesis and tail bud formation. However, when the epiboly delay becomes severe with higher dose of EtOH (2% and 3%), the delay and abnormalities were not fully restored causing stop of epiboly, incomplete gastrulation and failure of the germ ring closure. Such severe phenotype tends to cause failure of the complete tail bud formation leading to a short and disorganised tail structure (e.g. Mourabit et al. 2014). It has been reported that ethanol affects the microtubule cytoskeleton in the yolk syncytial layer resulting in the suppression of microtubule filament production, which is crucial for epiboly migration (Sarmah et al., 2020, Takesono et al. 2012) therefore such molecular mechanism might be the cause of the epiboly delay.

According to Blader and Strahle, 2.4% ethanol exposure at the gastrula stage inhibited the extension of the axial mesoderm, resulting in the failure of optical field separation and cyclopia (Blader and Strahle, 1998). Besides Tg(h2a:gfp), we have also used Tg(gsc:gfp) fish line for multiwell live imaging. In this line, GFP is expressed in the axial mesoderm including prechordal plate and notochord and is suitable for measuring the extension movement of these tissues. The data shows dose dependent delay of the extension and failure of the prechordal plate reaching to the animal pole. As seen in the delay of epiboly, the delay of axial mesoderm extension with mild dose (1%) seems restored at the end of gastrula to early somite stage, therefore the body size and morphology at 24 and 48hpf in 1% EtOH is similar to the control. Consistent with it, eye size and eye separation is relatively normal at 1% EtOH but in contrast with higher dose of EtOH, eye size is reduced, cyclopia is induced and such phenotype seems correlated with the delay of the failure of the prechordal plate reaching to the animal pole.

Although cell movement may explain many phenotypes of EtOH exposure, it is also important to examine gene expression as such change can also alter cell movement, cell differentiation, cell proliferation and consequently control tissue development and embryo morphogenesis. We have tested markers for three germ layers, endoderm, mesoderm and ectoderm. Overall pattern was that with 1% EtOH, gene expression was not suppressed except *hoxb1b*. But with 2% EtOH, gene expression was mostly suppressed except *ntl* and *gsc* which were less affected in terms of gene expression level. Cell movement including epiboly, convergent-extension movement were all highly suppressed at 2% EtOH seen in all germ layers markers including enoderm *(Sox17*), axial mesoderm (*gsc*) (Fig.4) and ectoderm (*p63, sox3*, Fig.5,6).

These data all together can highlight the change of the gene expression level and gene expression pattern of each gene. Overall, reduction and/or delay of gene expression occurred in all germ layers. This suggests that transcription process in general is possibly a target of the alcohol toxicity and the efficiency of the production of mRNAs is reduced.

To explain the microcephaly, reduction of the head neural ectoderm markers, *otx2* and *cyp26a1* seems important (Fig. 5). The area to be specified as head neural ectoderm (anterior neural ectoderm) is defined by the balance of Bmp, Bmp-angonists (e.g. Chordin) and posteriorizing activities (Wnt, Fgf and Retionic acid) (Kudoh et al. 2002, Kudoh et al. 2004). Therefore it is possible that these signalling activity is also compromised in the EtOH treated embryos, and consequently head neural induction is reduced and/or delayed.

*Sox3* is broadly expressed in the neural ectoderm and its expression showed a particularly complex pattern of response to EtOH. At 8hpf (80% epiboly for control), control embryos showed two expression domains, anterior neural ectoderm (animal-dorsal spot) and posterior neural ectoderm (band close to the blastoderm margin), however, embryos with 2% EtOH showed higher gene expression at this stage broadly spreading in the ectoderm without having two distinct expression domains (Fig. 5). However, by carefully examining earlier and later stages with and without EtOH, we found that the first wave of gene expression occurs from late blastula to early gastrula with broad ectodermal expression and the second wave of *sox3* expression occurs at mid to late gastrula stage in the two domains (anterior and posterior). By seeing these two steps of the *sox3* gene expression we concluded that the first wave of *sox3* expression is delayed by EtOH and the 2^nd^ wave of the *sox3* expression is strongly reduced with 2% ethanol.

Among all marker genes tested, the most highly suppressed gene by EtOH was *hoxb1b*, posterior neural ectoderm (prospective hindbrain and spinal cord) marker gene (Fig. 7). All other genes were only faintly reduced at 1% ethanol, but only this gene was highly reduced even at 1% EtOH and fully downregulated at 2% EtOH. The strong reduction of the *hoxb1b* expression seems to be caused by two mechanisms. Firstly, *hoxb1b* is induced by retinoic acid which is generated and secreted from the involuting paraxial mesoderm (Kudoh et al. 2002, Cruz et al. 2010). If involution of paraxial mesoderm is delayed, *hoxb1b* expression domain would become narrower and expression would become weaker. Secondly, In most of genes transcription event seems lower in the EtOH-treated embryos. Therefore in case of *hoxb1b*, these two effects by EtOH, cell involution movement and signalling, and general reduction of transcription may have caused additive suppression of the gene expression.

**Figure 7.**
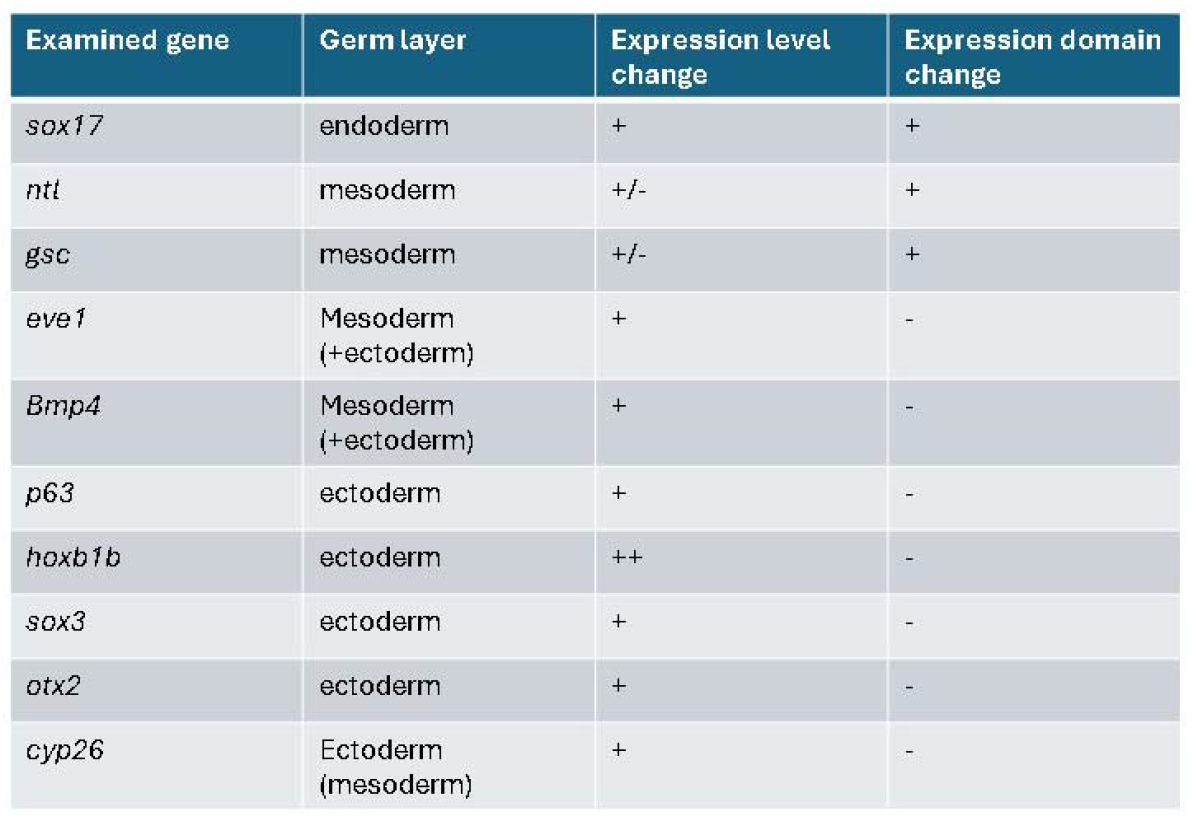
Summary of the in situ staining result. All tested genes except *ntl* and *gsc* showed clear suppression or delay of expression at 2% EthOH. With 1% EtOH, only the *hoxb1b* showed a clear reduction of the expression. In terms of expression domain, only the *sox17, ntl* and *gsc* expression domains showed clear alteration in morphology.

*Ntl* and *gsc* were the most unaffected by EtOH in terms of gene expression level (Fig. 7). The resistance to EtOH for these genes may be due to the fact that these are the earliest marker genes being expressed from mid blastula stage. All other markers are only expressed at late blastula to early gastrula stages. *Ntl* and *gsc* are induced by *Sqt* nodal signalling pathway (Kudoh et al. 2001) and signalling molecules of such pathway is already equipped from maternal storage in the egg. Therefore it is possible that the signalling was not largely affected.

Overall, our data revealed that all cell movements that we tested (convergence, extension, epiboly) are reduced by EtOH, and all genes expression are also reduced at some extent in a dose dependent manner. Some gene expression are more severely affected by EtOH possible due to the combinatorial effects of transcription regulation and indirect consequence of cell movement and signalling.

EtOH exposure in the first twenty-four hours of zebrafish development (including gastrulation phases) is equivalent to EtOH exposure in the third week of human pregnancy (Liu et al., 2012; Xu et al., 2010). It is highly conceivable that exposure of human foetus to EtOH would cause similar effect on both cell movement and gene expression in human foetus. Therefore the measurement of the epiboly and other cell movement in the small fish model can become a very easy measurement of the toxicity of EtOH. Our multi-well automated imaging assay system for examining the progress of epiboly is a very quick and high through put assessment for it. In this system, embryos are randomly oriented but by calculating the angle of the blastoderm margin (methods section), we were able to measure the epiboly with sufficient accuracy. To ensure our measurement is accurate enough, we have omitted some samples which orientation was wrongly angled (animal pole or vegetal pole facing to the camera). The large n number in the 96 well system further supported the accuracy of the live epiboly measurement. We are currently developing an automated software to measure epiboly using the fluorescent embryo and this would further enhance the efficiency of the testing of the EtOH toxicity in the fish embryos. A reliable and quantifiable system to determine the effect of ethanol on gastrulation would also allow the quick and efficient screening and assessment of drugs and nutrients that can reduce the toxicity of EtOH using this automated imaging system.

## Acknowledgement

AAA is funded by PSAU in Saudi Arabia. TK is funded by NC3R grant, NC/X001121/1. We would like to thank the Aquatic Resource Centre staff for maintaining the WT and TG zebrafish lines. We also thank to Anke Lange and John Dawdle for support.

## Notes

### Competing Interest Statement

The authors have declared no competing interest.

## References

Alsakran, A., and Kudoh, T. (2021). Zebrafish as a Model for Fetal Alcohol Spectrum Disorders. Front Pharmacol 12. doi: 10.3389/FPHAR.2021.721924.

Arenzana, F. J., Carvan, M. J., Aijón, J., Sánchez-González, R., Arévalo, R., and Porteros, A. (2006). Teratogenic effects of ethanol exposure on zebrafish visual system development. Neurotoxicol Teratol 28, 342–348. doi: 10.1016/J.NTT.2006.02.001.

Bilotta, J., Barnett, J. A., Hancock, L., and Saszik, S. (2004). Ethanol exposure alters zebrafish development: a novel model of fetal alcohol syndrome. Neurotoxicol Teratol 26, 737–743. doi: 10.1016/J.NTT.2004.06.011.

Blader, P., and Strähle, U. (1998). Ethanol impairs migration of the prechordal plate in the zebrafish embryo. Dev Biol 201, 185–201. doi: 10.1006/DBIO.1998.8995.

Carvan, M. J., Loucks, E., Weber, D. N., and Williams, F. E. (2004). Ethanol effects on the developing zebrafish: neurobehavior and skeletal morphogenesis. Neurotoxicol Teratol 26, 757–768. doi: 10.1016/J.NTT.2004.06.016.

Chahardehi, A. M., Arsad, H., and Lim, V. (2020). Zebrafish as a Successful Animal Model for Screening Toxicity of Medicinal Plants. Plants (Basel) 9, 1–35. doi: 10.3390/PLANTS9101345.

Finch, E., Cruz, C., Sloman, K. A., and Kudoh, T. (2010). Heterochrony in the germ ring closure and tail bud formation in embryonic development of rainbow trout (Oncorhynchus mykiss). J Exp Zool B Mol Dev Evol 314, 187–195. doi: 10.1002/JEZ.B.21325.

Fu, J., Han, N., Jiao, J., and Shi, G. (2021). Effects of Embryonic Exposure to Ethanol on Zebrafish Survival, Growth Pattern, Locomotor Activity and Retinal Development. Altern Ther Health Med 27, 120–128.

Iwamatsu, T. (2004). Stages of normal development in the medaka Oryzias latipes. Mech Dev 121, 605–618. doi: 10.1016/j.mod.2004.03.012.

Joya, X., Garcia-Algar, O., Vall, O., and Pujades, C. (2014). Transient exposure to ethanol during zebrafish embryogenesis results in defects in neuronal differentiation: an alternative model system to study FASD. PLoS One 9. doi: 10.1371/JOURNAL.PONE.0112851.

Kudoh T, Dawid IB. (2001) Role of the iroquois3 homeobox gene in organizer formation. Proc Natl Acad Sci U S A. 98:7852–7. doi: 10.1073/pnas.141224098.

Kudoh T, Wilson SW, Dawid IB. (2002) Distinct roles for Fgf, Wnt and retinoic acid in posteriorizing the neural ectoderm. Development. 129:4335–46. doi: 10.1242/dev.129.18.4335.

Kudoh, T., Concha, M. L., Houart, C., Dawid, I. B., and Wilson, S. W. (2004). Combinatorial Fgf and Bmp signalling patterns the gastrula ectoderm into prospective neural and epidermal domains. Development 131, 3581. doi: 10.1242/DEV.01227.

Lee O, Takesono A, Tada M, Tyler CR, Kudoh T. (2012). Biosensor zebrafish provide new insights into potential health effects of environmental estrogens. Environ Health Perspect. 120:990–6. doi: 10.1289/ehp.1104433.

Liu, J., and Lewis, G. (2014). Environmental Toxicity and Poor Cognitive Outcomes in Children and Adults. J Environ Health 76, 130. Available at: /pmc/articles/PMC4247328/.

Marrs, J. A., Clendenon, S. G., Ratcliffe, D. R., Fielding, S. M., Liu, Q., and Bosron, W. F. (2010). Zebrafish fetal alcohol syndrome model: effects of ethanol are rescued by retinoic acid supplement. Alcohol 44, 707–715. doi: 10.1016/J.ALCOHOL.2009.03.004.

Mourabit, S., Moles, M. W., Smith, E., Van Aerle, R., and Kudoh, T. (2014). Bmp suppression in mangrove killifish embryos causes a split in the body axis. PLoS One 9. doi: 10.1371/JOURNAL.PONE.0084786.

Muralidharan, P., Sarmah, S., Zhou, F. C., and Marrs, J. A. (2013). Fetal Alcohol Spectrum Disorder (FASD) Associated Neural Defects: Complex Mechanisms and Potential Therapeutic Targets. Brain Sci 3, 964. doi: 10.3390/BRAINSCI3020964.

Mourabit S, Fitzgerald JA, Ellis RP, Takesono A, Porteus CS, Trznadel M, Metz J, Winter MJ, Kudoh T, Tyler CR. (2019) New insights into organ-specific oxidative stress mechanisms using a novel biosensor zebrafish. Environ Int.133:105138. doi: 10.1016/j.envint.2019.105138.

Pinheiro-da-Silva, J., and Luchiari, A. C. (2021). Embryonic ethanol exposure on zebrafish early development. Brain Behav 11. doi: 10.1002/BRB3.2062.

Popova, S., Lange, S., Shield, K., Burd, L., and Rehm, J. (2019). Prevalence of fetal alcohol spectrum disorder among special subpopulations: a systematic review and meta-analysis. Addiction (Abingdon, England) 114, 1150. doi: 10.1111/ADD.14598.

Sarmah, S., Muralidharan, P., Curtis, C. L., McClintick, J. N., Buente, B. B., Holdgrafer, D. J., et al. (2013). Ethanol exposure disrupts extraembryonic microtubule cytoskeleton and embryonic blastomere cell adhesion, producing epiboly and gastrulation defects. Biol Open 2, 1013–1021. doi: 10.1242/BIO.20135546.

Sarmah, S., Srivastava, R., McClintick, J. N., Janga, S. C., Edenberg, H. J., and Marrs, J. A. (2020). Embryonic ethanol exposure alters expression of sox2 and other early transcripts in zebrafish, producing gastrulation defects. Sci Rep 10. doi: 10.1038/S41598-020-59043-X.

Soares, A. R., Pereira, P. M., Ferreira, V., Reverendo, M., Simóes, J., Bezerra, A. R., et al. (2012). Ethanol exposure induces upregulation of specific microRNAs in zebrafish embryos. Toxicol Sci 127, 18–28. doi: 10.1093/TOXSCI/KFS068.

Sylvain, N. J., Brewster, D. L., and Ali, D. W. (2010). Zebrafish embryos exposed to alcohol undergo abnormal development of motor neurons and muscle fibers. Neurotoxicol Teratol 32, 472–480. doi: 10.1016/J.NTT.2010.03.001.

Takesono A, Moger J, Farooq S, Cartwright E, Dawid IB, Wilson SW, Kudoh T. (2012) Solute carrier family 3 member 2 (Slc3a2) controls yolk syncytial layer (YSL) formation by regulating microtubule networks in the zebrafish embryo. Proc Natl Acad Sci U S A. 28;109:3371–6. doi: 10.1073/pnas.1200642109.

Wilhelm, C. J., and Guizzetti, M. (2016). Fetal alcohol spectrum disorders: An overview from the glia perspective. Front Integr Neurosci 9, 170319. doi: 10.3389/FNINT.2015.00065/BIBTEX.

Xu, T., Zhao, J., Hu, P., Dong, Z., Li, J., Zhang, H., et al. (2014). Pentachlorophenol exposure causes Warburg-like effects in zebrafish embryos at gastrulation stage. Toxicol Appl Pharmacol 277, 183–191. doi: 10.1016/J.TAAP.2014.03.004.

